# Ultrasensitive detection of neuronal DNA using genome-wide non-CG methylation patterns

**DOI:** 10.64898/2026.10.29.761818

**Authors:** Netanel Loyfer, Amit Horn, Shmuel Hart, Jonathan Rosenski, Sheina Piyanzin, Judith Magenheim, Ofra Sabag, Adi Avniel Aran, Yshia Langer, Natalie Barsoum, Guy Rosenthal, Moshe Gat, Kim Olthoff, Abraham Shaked, Ruth Shemer, Benjamin Glaser, Tommy Kaplan, Yuval Dor

**Affiliations:** Department of Developmental Biology and Cancer Research, The Institute for Medical Research Israel-Canada, Faculty of Medicine, The Hebrew University of Jerusalem, Jerusalem, Israel; School of Computer Science and Engineering, The Hebrew University of Jerusalem, Jerusalem, Israel; Section of Pediatric Critical Care, Hadassah Medical Center, Faculty of Medicine, The Hebrew University of Jerusalem, Israel; Department of Neurosurgery, Hadassah Medical Center, Faculty of Medicine, The Hebrew University of Jerusalem, Israel; Department of Hematology, Hadassah Medical Center, Faculty of Medicine, The Hebrew University of Jerusalem, Israel; Penn Transplant Institute, Department of Surgery, Perelman School of Medicine, University of Pennsylvania, Philadelphia, PA, USA; Endocrinology and Metabolism Service, Hadassah Medical Center, Faculty of Medicine, The Hebrew University of Jerusalem, Israel; Center for Computational Medicine, The Hebrew University of Jerusalem, Jerusalem, Israel; Barts Cancer Institute, Queen Mary University of London, London, UK; Centre for Epigenetics, Queen Mary University of London, London, UK; Prima Mente, London, UK

## Abstract

Methylation-based liquid biopsies can identify the tissue origins of circulating cell-free DNA (cfDNA). A major constraint on the sensitivity of these assays is the limited number of cell-type-specific CpG markers. Here we exploit the high abundance of non-CpG (CpH) methylation in neurons to enable ultrasensitive detection of neuronal DNA. We show that neurons carry 100-fold higher CpH methylation than other tissues, in which apparent CpH methylation largely reflects experimental artifacts. Using a computational criterion to identify cfDNA molecules carrying clusters of methylated CpA sites, we detect neuronal DNA at proportions as low as 20 parts per million. Analysis of 231 samples from 163 individuals revealed minimal neuronal cfDNA in the plasma of healthy subjects; elevated levels in patients with measles or acute neurological conditions; strikingly high levels in brain-dead donors, indicating extensive ongoing neuronal death; and the highest levels in the cerebrospinal fluid of patients in a neurological intensive care unit. These findings establish genome-wide, fragment-level analysis of non-CpG methylation as an ultrasensitive approach for blood-based monitoring of brain injury and disease.

## Main text

Maximization of assay sensitivity is a central challenge of liquid biopsies. In the context of cancer, early detection requires the ability to detect circulating tumor DNA when comprising less than 0.1% of the total^1^. Recent studies have revealed that this task is feasible if a large number of independent specific markers exists for the tissue of interest. For example, early detection of nasopharyngeal cancer, a tumor driven by EBV infection, was facilitated by the realization that each dying tumor cell releases to plasma 500 copies of a specific viral sequence^2^. More recently, ultrasensitive detection of circulating tumor DNA was demonstrated using hybrid capture panels of thousands of somatic mutations, deep sequencing of plasma cfDNA and methods for sequencing error correction^3–5^.

Epigenetic liquid biopsies based on DNA methylation patterns face a more fundamental limitation. Unlike somatic mutations or viral sequences, only dozens to a few hundred CpG loci exhibit methylation patterns that are truly unique to a given cell type^6^. As a result, methylation-based assays lack the degree of parallelism required for ultrasensitive detection. This limitation is particularly acute for brain-derived cfDNA, which is present at an extremely low concentration in plasma and may be preferentially released into the cerebrospinal fluid (CSF) rather than blood^7–10^. Consistent with this constraint, brain tumors are excluded from current methylation-based multi-cancer early detection assays such as GRAIL’s Galleri^11^, although plasma cfDNA methylome profiling has been used to discriminate between intracranial tumor types^12^.

Here we address this limitation by exploiting a fundamentally different class of epigenetic signal. The DNA of neurons is heavily methylated in non-CpG sites, compared with minimal levels of such methylation in other somatic tissues. Neurons methylate some cytosines followed by adenosine (CpA), and to a lesser extent cytosines followed by thymidine (CpT) or cytosine (CpC)^13–16^. While conserved in evolution, the underlying biology of this phenomenon is not fully understood^17,18^. Non-CpG (often termed CpH) methylation is thought to be mediated by the *de novo* DNA methyltransferase DNMT3A^19^, and to occur in genomic loci that are transcriptionally repressed in neurons^13^. CpH methylation accumulates postnatally, after neurons exit the cell division cycle^14,20,21^, consistent with the asymmetric nature of CpH sites, which leads to dilution with every division. In principle, the abundance of neuronal CpH methylation suggests a powerful and massively parallel biomarker for neuronal DNA. Neuronal CpH methylation has previously been proposed as a marker for brain-derived cfDNA, using targeted bisulfite sequencing of a small number of loci^22–24^. Such targeted panels, however, capture only a minute fraction of the millions of methylated CpH sites in the neuronal genome.

A major confounder in this approach is the reported presence of CpH methylation in 1-3% of non-neuronal DNA^15^. We show that this apparent background largely reflects technical under-conversion of unmethylated cytosines to uracils prior to sequencing, and that accounting for this artifact reveals a 100-fold contrast between neuronal and non-neuronal CpA methylation. Correcting for this background enables the use of genome-wide CpH methylation as an extremely specific and sensitive biomarker for neuronal DNA in mixtures, including the plasma of patients with brain pathologies.

### Global abundance of non-CpG methylation in humans

We first asked whether the global abundance of CpH methylation in neurons is sufficient to support ultrasensitive detection of neuronal DNA. To this end, we calculated genome-wide non-CpG methylation levels in multiple cell types, focusing on neurons and on the major contributors to cfDNA (hematopoietic cells, hepatocytes, and vascular endothelial cells)^6^, using a combination of public and in-house methylome datasets. In agreement with previous analyses, we found that bulk brain tissue exhibits substantial non-CpG methylation, with 3.4% of CpA sites methylated, as well as 1.3% of CpT sites and 0.75% of CpC sites (**Figure 1a**). Breaking down to constituent brain cell types, we found that in sorted cortical neurons, 9.8% of CpA sites were methylated, as well as 3.2% of CpT sites and 0.7% of CpC sites. Oligodendrocytes exhibited much lower methylation levels (1.5%, 0.5% and 0.25% for CpA, CpT and CpC, respectively). In all other tissues, 0.5-1% of non-CpG sites were methylated, in agreement with previous reports^15^.

**Figure 1.**
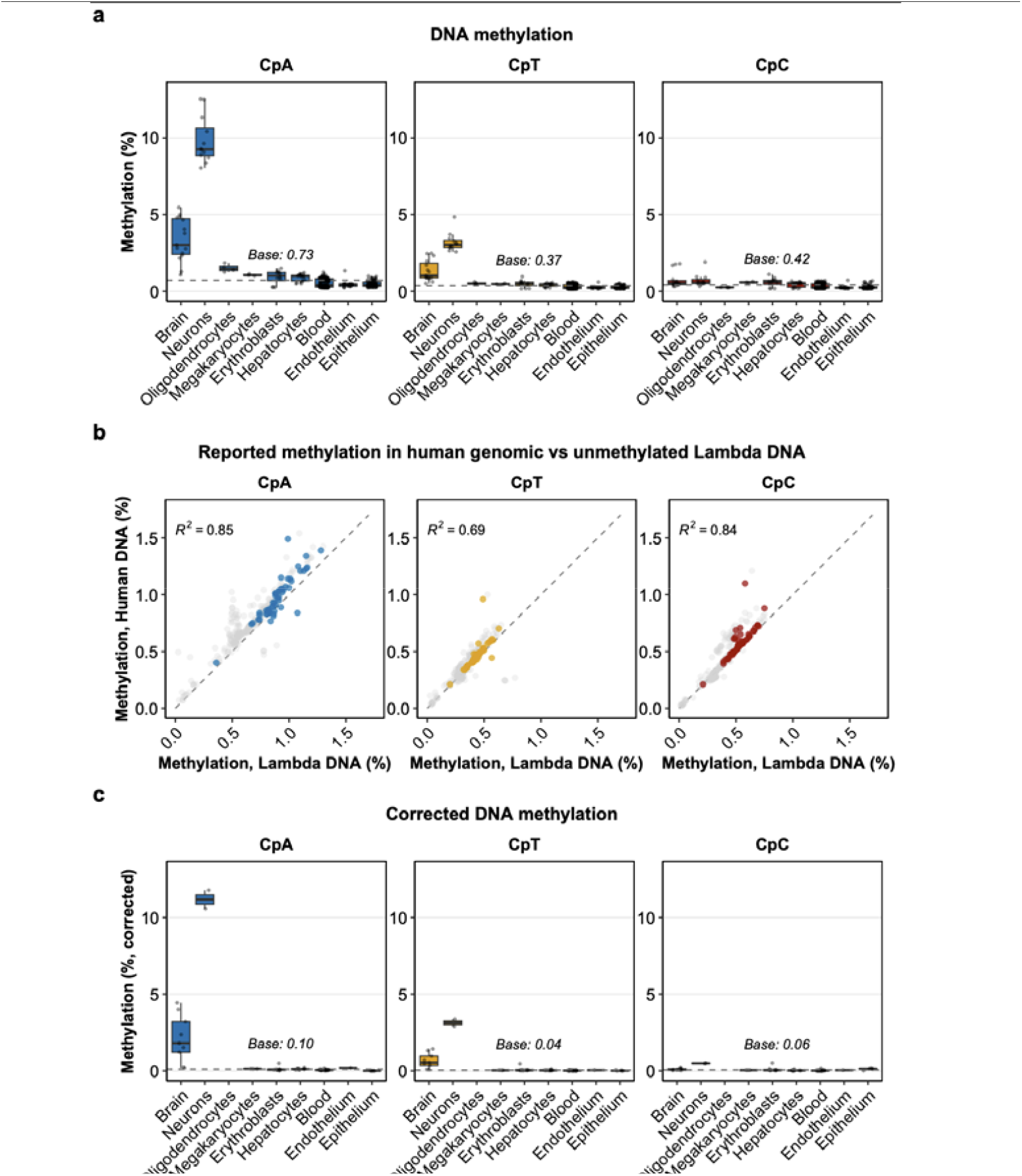
Global CpH methylation levels across human cell types and correction of technical background. **(a)** Global methylation percentages for CpA (blue), CpT (orange), and CpC (red) contexts across bulk brain tissue and sorted cell types. Box plots indicate the median (center line), interquartile range (IQR), and 1.5×IQR (whiskers). Sample sizes (n) are: bulk brain (n=7), neurons (n=12), oligodendrocytes (n=4), megakaryocytes (n=3), erythroblasts (n=12), hepatocytes (n=17), blood cells (n=167), endothelial cells (n=21), and epithelial cells (n=88). The horizontal dashed line (“Base”) indicates the mean CpH methylation level across all non-brain cell types. **(b)** Correlation between apparent CpH methylation measured in non-brain human DNA and spiked-in unmethylated λ phage DNA. Each point represents a single sample, including a subset of samples from a (n=53, colored) and additional bulk tissue, sorted cell types, and cfDNA samples (n=225, gray) from published and in-house datasets. Dashed lines indicate the line of identity (y=x). Pearson’s R^2^ values are shown for each CpH context. Brain-derived samples are excluded here because their high biological CpH methylation obscures the low-level technical background observed in λ DNA. **(c)** CpH methylation levels after correction for technical background in the subset of samples from **a** that include λ DNA spike-ins. Corrected values were obtained by subtracting the sample-specific λ methylation (technical background) from the corresponding human genomic CpH methylation. The higher median CpH methylation observed in neurons in **c** relative to **a** is due to the specific subset of samples containing spike-ins. Box plot elements are defined as in **a**.

The observation that nearly 10% of CpA sites are methylated in neurons suggests an unusually dense epigenetic signal and potentially an extremely sensitive biomarker. In fact, most of the ~18 million nucleosome-size cfDNA fragments released from each dying neuron (166bp-long, containing a median of 11 CpA sites) are expected to carry an identifiable covalent mark. However, the presence of 1% CpA methylation in non-neurons would provide a signal-to-noise ratio too low for utility as a biomarker for presence of rare neuronal DNA in complex mixtures.

### Apparent non-neuronal CpH methylation largely reflects technical artifacts

What are the origins of the apparent background CpH methylation in non-neuronal tissues? Most methods for detection of cytosine methylation, either chemical (bisulfite)^25^ or enzymatic (protective oxidation of methylated cytosines by TET followed by Apobec-mediated deamination)^26^ rely on selective deamination of unmethylated cytosines to uracils, while methylated cytosines are protected. Following PCR amplification and sequencing, unmethylated cytosines are read as thymidines and methylated cytosines are read as cytosines. The reaction of deamination is not perfect in either method, and a certain proportion of unmethylated cytosines (typically close to 1%) fails to convert to uracils. We therefore hypothesized that much of the CpH methylation observed in non-neuronal tissues reflects failed deamination rather than bona fide non-CpG methylation. To test this hypothesis, we analyzed unmethylated λ phage DNA, which is often used as an internal spike-in control. Consistent with a technical origin, the extent of CpH sites read as methylated in λ DNA correlated strongly with the apparent CpH methylation levels measured in non-neuronal human genomic DNA (**Figure 1b**). This correlation was observed across diverse tissues, cell types, and cfDNA samples, and across all non-CpG contexts (CpA, CpT and CpC), indicating that the background signal scales with sample-specific conversion efficiency. Brain-derived samples were excluded from this analysis, as their high endogenous CpH methylation obscures the low-level technical background.

This relationship enabled direct estimation of true CpH methylation in each sample, by subtracting the sample-specific artifactual λ-derived CpH signal from the corresponding genomic CpH measurements. Following this correction, CpH methylation in non-neuronal tissues was reduced to ~0.1% (**Figure 1c)**. In neurons, however, CpH methylation remained high, with corrected levels of 11% for CpA, 3.3% for CpT, and 0.5% for CpC. Thus, accounting for technical under-conversion reveals an effective signal-to-noise ratio of 100:1 between CpA methylation in neurons and non-neuronal tissues.

### Multiple methylated CpAs distinguish neuronal DNA fragments

Having established that true CpH methylation is abundant in neurons and minimal in other tissues, we next sought to develop a molecular-level classifier that could reliably distinguish neuron-derived DNA fragments from the vast background of blood cfDNA. To this end, we considered the distinct topologies of biological methylation and technical noise that arises from the chemistry of experimental deamination. CpG methylation typically exhibits a regional organization, with adjacent sites showing concordant patterns due to processive enzymatic activity^27,28^. We hypothesized that authentic CpH methylation in neurons would exhibit similar local clustering, whereas technical artifacts such as the failed bisulfite conversion of unmethylated cytosines would occur as stochastic, independent events (**Figure 2a**). Selected genomic loci harboring methylated CpA sites (mCpA) in neurons showed a pattern of methylation in neurons and blood that was consistent with this idea (**Figure 2b**).

**Figure 2.**
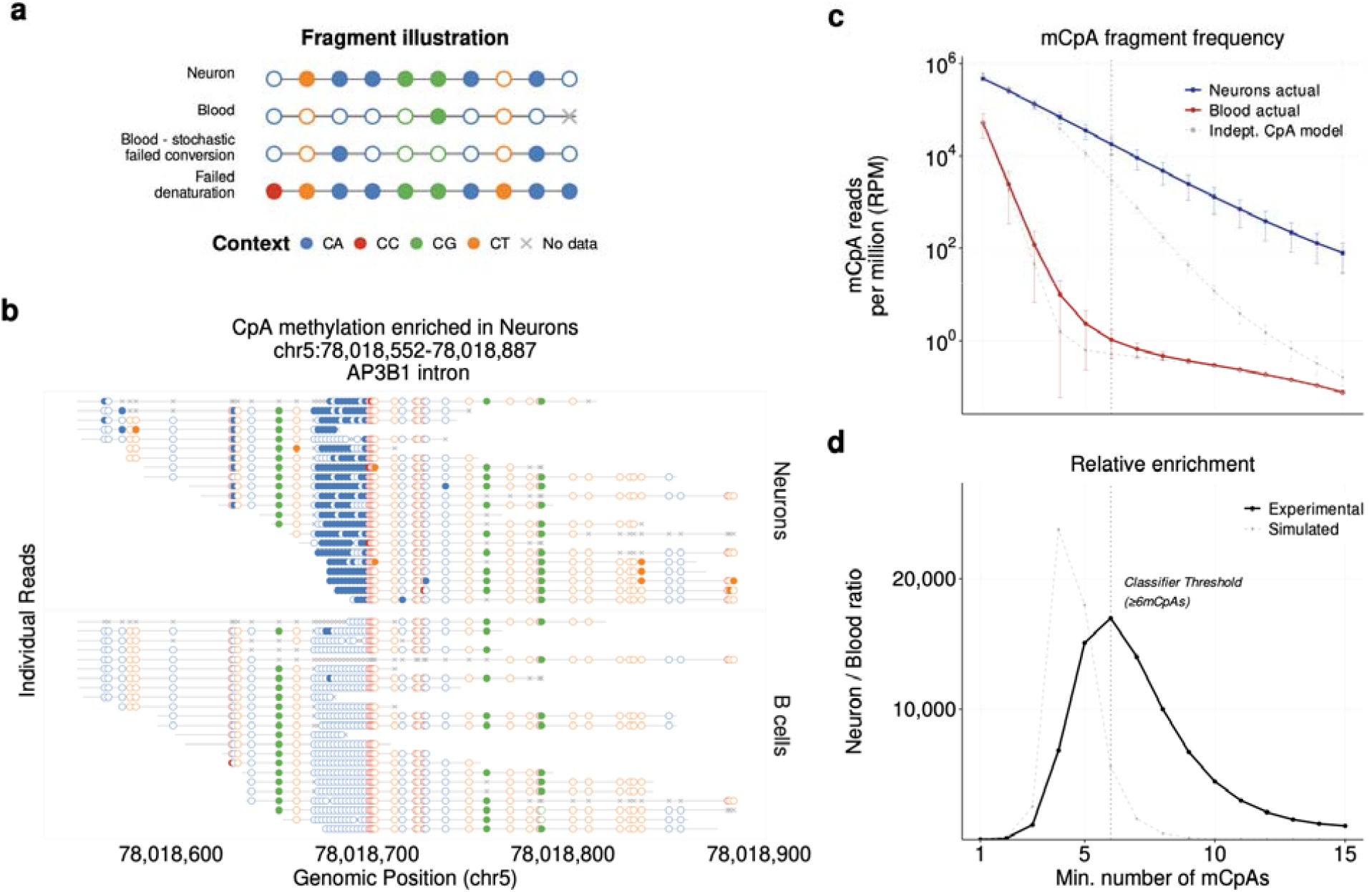
Fragment-level discrimination of neuronal DNA by clustered methylated CpAs. **(a)** Schematic illustrating presumed fragment-level DNA methylation patterns in neurons versus blood, and common technical artifacts. Authentic neuronal CpH methylation typically appears in clustered blocks, whereas artifacts arising from stochastic bisulfite conversion failure appear as isolated events. Molecules resulting from failed denaturation (incomplete conversion) mimic fully methylated reads. Empty and filled circles represent unmethylated and methylated cytosines, respectively. **(b)** Representative sequencing reads from neuronal genomic DNA (top) and blood B cells (bottom) at chr5:78,018,675-78,018,695, a locus exhibiting high CpA methylation in neurons. Top strand reads. **(c)** Number of reads per million (RPM) containing ≥k methylated CpA sites (mCpAs) in neuronal genomic DNA (blue, n=12 samples) and blood cell types (red, n=164 samples). Solid lines show experimental data; dashed lines show expectations from a generative null model assuming independent CpA methylation probabilities. Error bars indicate s.d. **(d)** The neuron-to-blood signal-to-noise ratio as a function of the minimum mCpA threshold. The experimental ratio (solid line) peaks at ≥6 mCpAs. In c and d, to exclude artifacts derived from failed denaturation, reads were additionally required to contain ≥2 unmethylated CpCs and no methylated CpCs.

To systematically test the clustering hypothesis, we compared experimental methylomes from neurons and blood to a generative null model that simulates CpA methylation as a random, independent process following a binomial distribution, parameterized by the observed global methylation rates (9.8% for neurons and 0.53% for blood). DNA fragments were scored by the number of mCpA sites they carried.

As expected under the null model, fragments with many mCpAs were rare in both neurons and blood. However, the experimental neuronal data deviated strongly from the expected, consistent with biological clustering. As the stringency threshold for methylated CpA increased (that is, requiring more methylated CpAs per fragment to classify a molecule as neuron-derived), the blood background decayed rapidly, closely matching the null prediction. In contrast, the neuronal signal decayed much more slowly, indicating an excess of fragments carrying multiple mCpAs beyond what would be expected by chance **(Figure 2c)**. For example, at a threshold of methylated CpA≥6 per read, the null model predicted only 2,944 reads per million (RPM) in neuronal DNA, whereas the experimental data yielded 17,943 RPM (after the denaturation filter described below), representing a six-fold enrichment driven by biological clustering of CpA methylation (**Figure 2c**). This divergence between model and experimental data underlies the discriminatory power of the classifier.

While the simulated signal-to-noise ratio (assuming no clustering of methylated CpA) peaked at ≥4 mCpAs and declined thereafter, the experimental neuron-to-blood ratio continued to increase with stringency up to ≥6 mCpAs, where neuronal DNA yielded 17,943 RPM and blood cells 0.53 RPM (Figure 2c-d). Stricter criteria reduced the neuronal read count without further increasing discrimination. We therefore selected the presence of at least 6 methylated CpA sites as an operational criterion for neuronal DNA.

Incomplete denaturation of DNA represents an additional, qualitatively distinct source of technical noise. Because bisulfite deamination requires single-stranded DNA, failed denaturation protects all cytosines from conversion, producing fragments that appear fully methylated (**Figure 2a**). To exclude such artifacts, we exploited the scarcity of authentic CpC methylation in neurons. Fragments were classified as neuron-derived only if they contained at least two unmethylated CpC sites and no methylated CpCs, providing an internal control for successful denaturation and bisulfite activity. At ≥6 mCpAs, this filter reduced the blood background ~70-fold (from 39 to 0.53 RPM) while retaining 78% of neuronal reads (from 23,071 to 17,943 RPM), raising the neuron-to-blood ratio from ~600:1 to ~34,000:1 (~17,000:1 with the pseudocount regularization used in **Figure 2d**; see Methods).

Together, these analyses establish a robust molecule-level classifier to distinguish neuronal from non-neuronal DNA molecules with high sensitivity and accuracy.

### Ultrasensitive detection of neuronal DNA in plasma

Having established a robust fragment-level classifier for neuronal DNA, we next evaluated its sensitivity in plasma-like mixtures. We performed *in silico* serial dilution experiments in which neuronal methylomes were computationally spiked into simulated cfDNA plasma methylomes composed of blood cell types, hepatocytes and vascular endothelial DNA in proportions typical of healthy plasma. Neuronal signal was estimated using one billion reads (corresponding to whole-methylome sequencing at 50x depth). Each simulation was repeated 20 times, by randomly subsampling the respective methylomes.

The classifier readily detected the presence of neuronal DNA in dilutions down to 1:100,000, and failed to detect neuronal DNA when comprising 1:1,000,000 of the mixture, relative to simulated healthy plasma (**Figure 3a**). This analytical sensitivity is in line with the theoretical performance of the classifier (**Figure 2**). For comparison, we analyzed the same simulated mixtures using neuron-specific CpG methylation markers^6^ (see Methods). This CpG-based analysis picked up neuronal DNA only when comprising at least 1:1,000 of the mixture, suggesting that CpH-based analysis is two orders of magnitude more sensitive than current state-of-the-art CpG-based analyses (**Figure 3b**).

**Figure 3.**
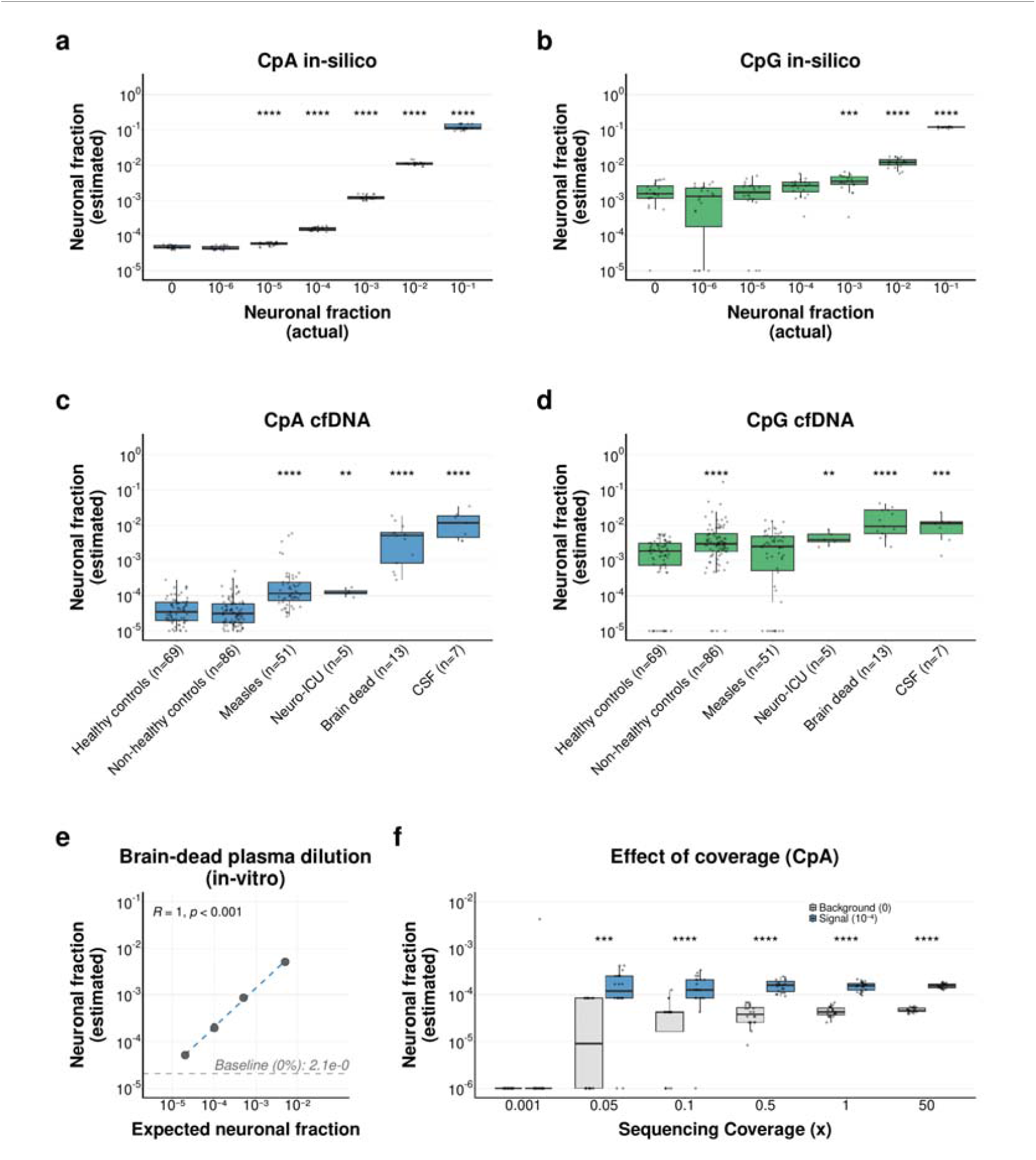
Ultrasensitive detection of neuronal DNA in simulated plasma and in patient samples. **(a-b)** *In silico* mixtures. Neuron-derived methylome was computationally spiked into simulated cfDNA plasma methylome samples with coverage of 50x. The simulated samples were composed of DNA from leukocytes, megakaryocytes, endothelial cells, erythroblasts, and hepatocytes, combined in ratios typical of healthy plasma. Analysis was performed using the criterion of ≥6 methylated CpA sites per sequence read **(a)**, or using deconvolution based on the neuronal signal in our CpG methylome atlas (**b**). **(c-d)** Neuronal cfDNA signal measured in plasma cfDNA from healthy individuals and patients (CSF, cerebrospinal fluid cfDNA; repeated samples from the same individuals are included for non-healthy controls, measles, neuro-ICU, brain-dead donors and CSF), using CpA methylation (**c**) or CpG methylation (**d**). **(e)** *In vitro* validation. Plasma from a brain-dead donor (measured neuronal fraction: 0.51%) was mixed into healthy control plasma at ratios of 1:10, 1:50, and 1:250. The x-axis represents the expected neuronal fraction calculated based on the 0.51% baseline. The regression (Pearson’s R = 1, rounded to two decimals; P < 0.001) is calculated on log-transformed data, excluding the 0% mixture. Horizontal dashed line represents the estimated neuronal fraction in healthy plasma (same sample into which brain-dead plasma was spiked in). **(f)** Limit of detection (LOD) analysis (*in silico*). Performance of the CpA method evaluated at varying sequencing coverages (0.001x to 50x), and when neuronal DNA comprises 0.01% or 0.001% of the total. Statistical significance is calculated relative to the 0% dilution (in panels a+b) or relative to healthy controls (in panels c+d). * P≤0.05; ** P≤0.01; *** P≤0.001; **** P≤0.0001 (Wilcoxon rank-sum test).

We then applied the assay to actual plasma samples sequenced at depths ranging from 4.4x to 165x (median 32x), converting the proportion of mCpA reads to estimates of neuronal DNA contribution based on the mCpA profile in isolated neurons (see Methods). Among 69 samples from healthy adults, the neuronal cfDNA signal comprised 0.0052% of the total (1:19,055, or 52 parts per million, **Figure 3c**), reflecting a combination of potential true neuronal DNA and baseline noise. A similar low-level signal was observed in 86 samples from 68 patients with pathologies not involving the brain **(Figure 3c)**. The highest value among these patients (0.05% neuronal signal in plasma) corresponded to a patient with multiple myeloma complicated by spinal cord compression necessitating irradiation, consistent with collateral neuronal damage.

Significantly elevated levels were observed in patients treated in a neurological intensive care unit (5 samples from 2 patients, 0.013%, P=7×10^−4^) and in children with severe measles, hospitalized in a pediatric critical care unit (51 samples from 20 patients, 0.048%, P=1×10^−10^), suggesting that neuronal death in these conditions releases detectable cfDNA to the bloodstream. Most strikingly, plasma samples from brain-dead donors (13 samples from 4 donors) exhibited markedly elevated concentrations of neuronal cfDNA (~0.55%, P=1×10^−8^), consistent with massive ongoing brain cell death (**Figure 3c**). As a positive control, cerebrospinal fluid (CSF) obtained from the same two neurological-ICU patients (7 samples) carried far higher levels of neuronal cfDNA (mean 1.4%, P=9×10^−10^), approximately 100-fold above their matched plasma (**Figure 3c**).

For comparison, CpG-based analysis showed a substantially broader signal distribution among patients without brain pathologies (22 of 86 above the range of healthy controls, versus 2 of 86 by CpA), and failed to separate 3 of 13 brain-dead samples from healthy controls (**Figure 3d**). This reduced specificity likely reflects confounding contributions of globally hypomethylated erythroblasts to cfDNA^6,29^, which limit the detection of tissue-specific contributions based on demethylated CpGs.

We took advantage of the high proportion of neuronal DNA detected in the plasma of brain-dead donors, and performed a more rigorous plasma-to-plasma spike-in wet experiment. Genome-wide analysis of mCpA revealed a distinct neuronal signal even when brain-dead plasma (from a donor with an estimated neuronal cfDNA contribution of 0.51%) was diluted 1:250 into normal plasma. Thus, the fraction of neuronal cfDNA that could be detected was 0.0020%, or 20 parts per million (**Figure 3e**). This suggests a real-life limit of detection (LOD) of 1:49,362 for detecting neuronal cfDNA in plasma, in line with theoretical considerations.

### Neuronal DNA detection using shallow sequencing

A hallmark feature of CpA methylation is its broad distribution across the genome. While this may limit the utility of targeted analysis based on deep sequencing of few selected loci, shallow whole-methylome sequencing may suffice to detect a signal, with important implications for deployment of a practical blood test. To test this idea, we assessed how sensitivity of detection was affected by sequencing depth. We reanalyzed the simulated neuronal spike-in data shown in **Figure 3a**, randomly selecting fewer sequenced reads that corresponded to lower coverage. We asked how reduced sequencing depth affected the ability to detect a neuronal signal. Strikingly, the analysis suggested that sequencing depth as low as 0.05x (i.e., one million reads) was sufficient to pick up a neuronal signal comprising 0.01% of the total (**Figure 3f**). The performance of shallow sequencing in this experiment suggests that it will be feasible to detect neuronal cfDNA using minute amounts of blood, and to use near-real time sequencing methods in clinical settings that require a short turnaround time.

## Discussion

A central insight emerging from recent advances in the liquid biopsy field is that ultrasensitive detection of cfDNA requires hundreds to thousands of independent biomarkers^2–5^. When such markers are examined at maximal depth (i.e., assessing all available copies in a plasma sample, approximately 1,000 genome equivalents per mL), this yields on the order of one million detection opportunities, enabling in principle sensitivity in the parts-per-million range. The performance of such an approach, however, depends crucially on maintaining an extremely low background. In mutation-based liquid biopsy assays, the dominant source of background noise arises from sequencing errors, necessitating the use of dedicated error-correction algorithms to suppress noise and achieve reliable detection.

Methylation-based epigenetic liquid biopsy enables detection of the tissue origins of circulating cell-free DNA and is increasingly used to identify pathological processes involving excessive cell death, including cancer, without relying on mutations as biomarkers. The principles governing assay performance are similar, in that maximal sensitivity is determined by the number of cell type-specific methylation markers assessed in parallel. This has been a major limitation, as methylation atlases have identified only dozens to a few hundreds of absolutely unique markers per cell type^6^. Specificity is further constrained by the efficiency of deamination, with artifacts resulting from both over- and under-conversion of cytosines (methylated or unmethylated, respectively) to uracils^30^. Sequencing errors also contribute to noise, but this is of lesser concern, as sequencing errors across multiple nucleotides in the same read are exceedingly rare.

Here we applied these principles to develop a biomarker for neuronal cfDNA. High sensitivity is enabled by the unique presence of millions of non-CpG methylated cytosines in neurons, while specificity is achieved by exploiting the block-like behavior of authentic DNA methylation in contrast to the random nature of failed conversion. Although this study focused on classical bisulfite-mediated methylation analysis, the same principles and challenges apply to the more recently developed enzyme-based methylation assays.

Our work extends an earlier proposal to use CpH methylation at selected loci for detecting brain-derived cfDNA^22–24^, in several respects. First, we show that much of the apparent CpH methylation in non-neuronal DNA reflects incomplete deamination rather than true methylation, and we define a fragment-level criterion that is robust to this artifact. Second, rather than relying on a predefined panel of loci, we score every sequenced cfDNA fragment across the genome for clustered methylated CpAs, thereby exploiting the millions of CpA sites methylated in neurons. Third, this genome-wide approach reaches an analytical sensitivity in the parts-per-million range, two orders of magnitude beyond CpG-based analysis, and in simulations allows detection with shallow sequencing.

The approach described here has the potential to form the basis of a new type of blood test for detection of neuronal cfDNA, providing a direct readout of neuronal cell death, an unmet need in many brain pathologies. Some applications are in the field of cancer, where the presence of neuronal cfDNA would signal the impact of cancer cells or therapy on genetically normal neurons. Additional applications include the detection of brain involvement in infectious (e.g., measles) and autoimmune diseases (e.g., multiple sclerosis), and the monitoring of brain damage in neurodegenerative diseases and following stroke or trauma. Notably, for some of these pathologies, we and others have already demonstrated the presence of brain-derived cfDNA, using lower-sensitivity assays based on CpG methylation markers^31–35^.

We note that there is substantial room for further enhancement of assay performance. For example, our current analysis does not take into account genomic context, such as loci with particularly high CpA methylation in neurons, which could further increase the signal-to-noise ratio. In addition, while our approach relies on whole-methylome sequencing, pulling down the most informative loci and sequencing these at a greater depth (potentially scoring all available molecules present in a blood sample) is expected to further increase assay sensitivity and specificity. Furthermore, computational approaches that consider additional features, such as fragment length and fragment ends, may increase performance further. Finally, the abundance of neuronal methylation markers suggests novel diagnostic applications, such as the use of much lower blood volume (e.g., a finger prick) or near-real-time analysis using nanopore sequencing, which has recently been applied to native methylation profiling of plasma cfDNA^36^.

In conclusion, the exceptional abundance of non-CpG methylation in neurons provides an opportunity for detection of neuronal DNA, at a sensitivity in the parts per million range, providing a powerful new avenue for monitoring brain injury and disease through blood-based assays.

## Methods

### Patient samples

Deidentified brain-dead donor samples were obtained from the BioTIP repository (Biorepository of the Transplant Institute at Penn), under approval by the University of Pennsylvania IRB. Samples from pediatric patients with measles and from patients in a neurological intensive care unit (both plasma and CSF) were collected at Hadassah Medical Center under institutional review board approval #0198-14-HMO (principal investigator: Prof. Benjamin Glaser). For all plasma samples, blood was collected in Streck cfDNA tubes (Streck) and processed within 7 days of collection. Blood was centrifuged at room temperature for 10 minutes at 1,600 × g, then 15 minutes at 4,122 × g. Plasma was aliquoted and stored at −80 °C until use.

### Extraction and processing of DNA

Genomic DNA was extracted using a Qiagen genomic DNA extraction kit. cfDNA was extracted from plasma using a Qiagen EZ2 liquid handling robot. We measured fragment size distributions using TapeStation, and quantified the concentration of cfDNA using Qubit. DNA was treated with bisulfite using a Zymo kit per manufacturer instructions, followed by whole-genome bisulfite sequencing library construction and sequencing on an Ultima Genomics UG 100 or an Illumina NovaSeq 6000 sequencer. Plasma and CSF samples were sequenced to a mean genome-wide coverage of 4.4x–165x (median 32x).

### Whole-genome bisulfite sequencing (WGBS) and data processing

For samples analyzed via the Ultima Genomics UG 100™ platform, read alignment and deduplication were performed using the proprietary Ultima Genomics Analysis Pipeline with its integrated, platform-specific mapper. Samples processed on the Illumina NovaSeq 6000 system were sequenced using a 2 × 150 bp paired-end strategy. Resulting raw Illumina FASTQ files were aligned to the human reference genome (hg38) utilizing bwa-meth^37^.

### Global DNA methylation analysis

To assess global DNA methylation across CpH dinucleotide contexts (CpA, CpT, and CpC) (**Figure 1a**), we calculated genome-wide methylation levels for both in-house generated libraries and publicly available datasets. The analysis was restricted to purified cell populations representing common contributors to the human cfDNA pool, as well as sorted brain-specific cell types and bulk brain tissue. Publicly available datasets were obtained from previous studies ^6,38,39^ and accessions are provided in the original publications.

### Background correction

For background calibration (**Figure 1b**), the correlation between measured methylation in human genomic DNA and spiked-in unmethylated λ phage DNA was evaluated. This analysis included the primary cohort (n=53) and an expansion set of 225 tissue, sorted-cell, buffy-coat and cell-free DNA methylomes (n=278 in total). Pearson’s R^2^ was calculated to evaluate the linearity of technical background noise across CpH contexts (CpA, CpT, and CpC). Brain-derived samples were excluded from the correlation analysis in **Figure 1b**, as high endogenous CpH methylation levels (biological signal) in these samples obscure the low-level technical background observed in the λ DNA. Background correction (**Figure 1c**) was performed by subtracting the sample-specific λ methylation percentage from the corresponding human genomic CpH methylation percentage for each context. Corrected values resulting in negative percentages were floored to zero.

### Generative modeling of independent methylation

To evaluate the distribution of methylated CpA sites under an assumption of independence, we developed a generative null model. The model utilized the total number of reads containing i CpA sites (where i ranges from 1 to 20) derived from the experimental data. For neurons and blood, we calculated a global methylation rate p (0.098 for neurons and 0.0053 for blood). For every read with i sites, the number of methylated sites j was simulated by drawing from a binomial distribution B(i, p), assuming that the methylation probability of each site is independent of its neighbors.

### Processing and signal-to-noise analysis

Both experimental and simulated data were processed to count the frequency of reads in neuron (n=12) and blood (n=164) samples containing exactly j methylated CpA sites out of i CpA sites. These counts were normalized to Reads Per Million (RPM). To model unaccounted for experimental noise and batch effect differences, and to mitigate artifacts arising from sampling zeros in the blood data when calculating ratios, we applied a regularization step by adding pseudo-counts totaling approximately 1 RPM, split uniformly across all 210 combinations of CpA sites and methylated CpAs (0.005 RPM each) in both cell types. We then aggregated these values to determine the cumulative abundance of reads with ≥k mCpA sites. The signal-to-noise ratio was calculated as the ratio of the cumulative RPM in neurons to the cumulative RPM in blood at each threshold k.

### Estimation of the neuronal cfDNA fraction

The neuronal cfDNA fraction was estimated as the proportion of reads carrying ≥6 methylated CpAs (after the CpC conversion filter) among all reads, multiplied by a calibration factor of 85. Because the probability that a read carries ≥6 methylated CpAs increases with read length, the factor was derived from nine sorted neuron methylomes, with each neuron’s rate weighted to the read-length distribution of healthy plasma cfDNA (mean, 11,825 reads with ≥6 methylated CpAs per million reads; 85 ≈ 10^6^/11,825), so that a pure neuronal sample with plasma-like fragment lengths is estimated at 1.

### Detection of neuronal DNA based on CpG markers

Cell-type deconvolution was performed using a modified version of the UXM method^6^, hereafter referred to as “binary deconvolution”. This approach utilizes a non-negative least squares (NNLS) regression framework applied to a reference atlas containing two primary categories: a neuronal signal and a consolidated background. The atlas incorporates the top 250 neuronal markers, which were identified using a modified version of the *find_markers* utility in wgbstools^40^. Specifically, a methylated block was defined as a neuronal marker if it contained ≥4 CpG sites and exhibited a difference in the proportion of unmethylated fragments >0.7 between neuronal and non-neuronal reference samples.

### Data sources and availability

We used data published in our methylome atlas^6^ and by others^38,39^. The previously published whole-genome bisulfite sequencing (WGBS) data utilized in this study have been previously deposited to the European Genome-phenome Archive (EGA) under study accession number: EGAS00001006791^6^. Due to participant privacy, raw sequencing data for the new samples are not publicly deposited but will be made available by the corresponding author following publication of the peer-reviewed article, upon reasonable request and for non-commercial research use.

## Code availability

The custom code used for data analysis in this study is based on a modified version of wgbstools^40^. The code for using a binary CpG deconvolution is available at https://github.com/nloyfer/UXM_deconv

## Acknowledgements

We thank Ariel Jaimovich, Gat Krieger, Eti Meiri, and Zohar Shipony for fruitful discussions.

## Ethics declarations

Competing interests: B.G., N.L., R.S., T.K., and Y.D. have filed patents on cell-free DNA methylation. N.L. is currently an employee of Prima Mente.

